# A haplotype-resolved draft genome of the European sardine (*Sardina pilchardus*)

**DOI:** 10.1101/441774

**Authors:** Bruno Louro, Gianluca De Moro, Carlos Garcia, Cymon J. Cox, Ana Veríssimo, Stephen J. Sabatino, António M. Santos, Adelino V. M. Canário

**Author notes:** authors contributed equally (Bruno Louro, Gianluca De Moro). Corresponding author: Adelino V. M. Canário.

## Abstract

**Background:** The European sardine (*Sardina pilchardus* Walbaum, 1792) has a high cultural and economic importance throughout its distribution. Monitoring studies of sardine populations report an alarming decrease in stocks due to overfishing and environmental change, which has resulted in historically low captures along the Iberian Atlantic coast. Consequently, there is an urgent need to better understand the causal factors of this continuing decrease in the sardine stock. Important biological and ecological features such as levels of population diversity, structure, and migratory patterns can be addressed with the development and use of genomics resources.

**Findings:** The sardine genome of a single female individual was sequenced using Illumina HiSeq X Ten 10X Genomics linked-reads generating 113.8 Gb of data. Three draft genomes were assembled: two haploid genomes with a total size of 935 Mbp (N50 103Kb) each, and a consensus genome with a total size of 950 Mbp (N50 97Kb). The genome completeness assessment captured 84% of Actinopterygii Benchmarking Universal Single-Copy Orthologs. To obtain a more complete analysis, the transcriptomes of eleven tissues were sequenced and used to aid the functional annotation of the genome, resulting in 40 777 genes predicted. Variant calling on nearly half of the haplotype genome resulted in the identification of more than 2.3 million phased SNPs with heterozygous loci.

**Conclusions:** A draft genome was obtained with the 10X Genomics linked-reads technology, despite a high level of sequence repeats and heterozygosity that are expected genome characteristics of a wild sardine. The reference sardine genome and respective variant data are a cornerstone resource of ongoing population genomics studies to be integrated into future sardine stock assessment modelling to better manage this valuable resource.

## Data description

### Background

The European sardine (*Sardina pilchardus* Walbaum, 1792) (NCBI:txid27697, Fishbase ID:1350) (Figure 1) is a small pelagic fish occurring in temperate boundary currents of the Northeast Atlantic down to Cape Verde off the west coast of Africa, and throughout the Mediterranean to the Black Sea [1]. Two subspecies are generally recognised: *Sardina pilchardus pilchardus* occupies the north-eastern Atlantic and the North Sea whereas *S. pilchardus sardina* occupies the Mediterranean and Black seas, and the North African coasts south to Cape Verde, with a contact zone near the Strait of Gibraltar [1, 2]. As with other members of the Clupeidae family (e.g. herring, *Clupea harengus*, Fishbase ID:24) and allis shad (*Alosa alosa*, NCBI: txid278164, Fishbase ID:101) [3], the sardine experiences strong population fluctuations in abundance, possibly reflecting environmental fluctuations, including climate change [4, 5].

**Figure 1.**
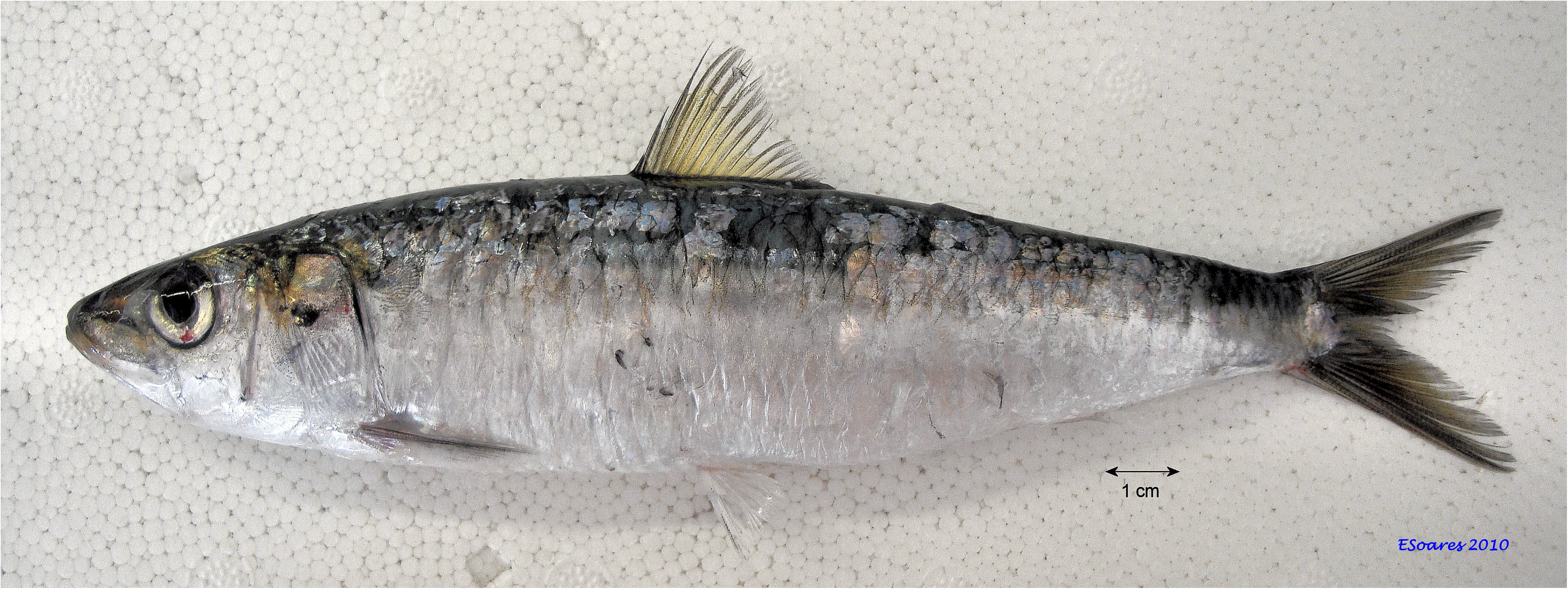
The European sardine, *Sardina pilchardus* (photo credit ©<u>Eduardo Soares, IPMA</u>)

The sardine is of major economic and social importance throughout its range with a reported commercial catch for 2016 of 72 183 tonnes in European waters [6]. In Portugal, the sardine is an iconic and culturally revered fish and plays a central role in tourist events, such as summer festivals, throughout the country. However, recent stock assessment data strongly suggests the Iberian sardine fisheries is under threat. A recent report by the International Council for the Exploration of the Sea [6] noted a sharp decrease in the Iberian Atlantic coast sardine stock and advised that catches in 2017 should be no more than 23 000 tonnes. The sardine fishery biomass has suffered from declining annual recruitment between 1978 and 2006, and more recently, it has fluctuated around historically low values indicating a high risk of collapse of the Iberian Atlantic stocks [6].

A number of sardine populations have been identified by morphometric methods, including as many as five populations in the north-eastern Atlantic (including the Azores), two off the Moroccan coast, and one in Senegalese waters [1, 7]. Each of these recognized sardine populations is subjected to specific climatic and oceanic conditions, mainly during larval development, which directly influence the recruitment of the sardine fisheries [4, 8, 9]. However, because of phenotypic plasticity, morphological traits are strongly influenced by environmental conditions and the underlying genetics that define those populations has proven elusive [10]. While the recognition of subspecies and localised populations might indicate significant genetic structure, the large population sizes and extensive migration of sardines are likely to increase gene flow and reduce population differences, suggesting, at its most extensive, a panmictic population with little genetic differentiation within the species’ range [11].

It is now well established that to fully understand the genetic basis of evolutionarily and ecologically significant traits, the gene and regulatory element composition of different individuals or populations needs to be assessed [see e.g., 12, 13]. Therefore, we provide a European sardine draft genome, providing the essential tool to assess the genetic structure of the sardine population(s) and for genetic studies of the life-history and ecological traits of this small pelagic fish, which will be instrumental for conservation and fisheries management.

### Genome sequencing

Sardines were caught during commercial fishing operations in the coastal waters off Olhão, Portugal, and maintained live at the experimental fish culture facilities (EPPO) of the Portuguese Institute for the Sea and Atmosphere (IPMA), Olhão, Portugal [14]. A single adult female was anesthetised with 2-phenoxyethanol (1:250 v/v), blood was collected in a heparinized syringe, and the fish euthanized by cervical section. Eleven tissues were dissected out – gill together with branchial arch, liver, spleen, ovary, midgut, white muscle, red muscle, kidney, head kidney, brain together with pituitary, and caudal fin (including skin, scales, bone and cartilage) – into RNA*later* (Sigma-Aldrich, USA) at room temperature followed by storage at –20□°C. Fish maintenance and sample collection were carried out in accordance with the guidelines of the European Union Council (86/609/EU) and Portuguese legislation for the use of laboratory animals from the Veterinary Medicines Directorate (DGAV), the Portuguese competent authority for the protection of animals, Ministry of Agriculture, Rural Development and Fisheries, Portugal (permit 010238 of 19/04/2016).

Total RNA was extracted using a total RNA purification kit (Maxwell^®^ 16 Total RNA Purification Kit, Promega) and digested twice with DNase (DNA-free kit, Ambion, UK). The total RNA samples where kept at −80°C until shipment to the RNAseq service provider Admera Health Co. (USA) which confirmed a RIN above 8 (Qubit Tapestation) upon arrival. The mRNA library preparation was performed with NEBNext^®^ Poly(A) mRNA Magnetic Isolation Module kit and NEBNext^®^ Ultra™ Directional RNA Library Prep kit for sequencing using Illumina HiSeq 4000 paired-end 150 bp cycle to generate about 596 million paired-end reads in total.

The genomic DNA (gDNA) was isolated from 20 μl of fresh blood using the DNeasy blood and tissue kit (Quiagen), followed by RNase treatment according to the manufacturer’s protocol. The integrity of the gDNA was confirmed using pulsed-field gel electrophoresis and showed fragment sizes largely above 50 kbp. The gDNA was stored at –20□°C before shipping to the service provider (Genome.one, Darlinghurst, Australia). Microfluidic partitioned gDNA libraries using the 10x Genomics Chromium System were made using 0.6 ng of gDNA input. Sequencing (150bp paired-end cycle) was performed in a single lane of the Illumina HiSeq X Ten instrument (Illumina, San Diego, CA, USA). Chromium library size range (580-850 bp) was determined with LabChip GX Touch (PerkinElmer) and library yield (6.5-40□M) by quantitative polymerase chain reaction.

### Genome size estimation

A total of 759 million paired-end reads were generated representing 113.8 Gb nucleotide sequences with 76.1% bases >= Q30. Raw reads were edited to trim 10X Genomics proprietary barcodes with a python script “filter_10xReads.py” [15] prior to kmer counting with Jellyfish v2.2.10 (Jellyfish, RRID:SCR_005491) [16]. Six hundred and seventy million edited reads (90.5 Gb) were used to obtain the frequency distribution of 23-mers. The histogram of the kmer counting distribution was plotted in GenomeScope v1.0.0 (Genoscope, RRID:SCR_002172) [17] (Figure 2) with maximum kmer coverage of 10 000 for estimation of genome size, heterozygosity and repeat content. The estimated sardine haploid genome size was 907 Mbp with a repeat content of 40.7% and a heterozygosity level of 1.43% represented in the first peak of the distribution. These high levels of heterozygosity and repeat content indicated a troublesome genome characteristic for *de novo* assembly.

**Figure 2.**
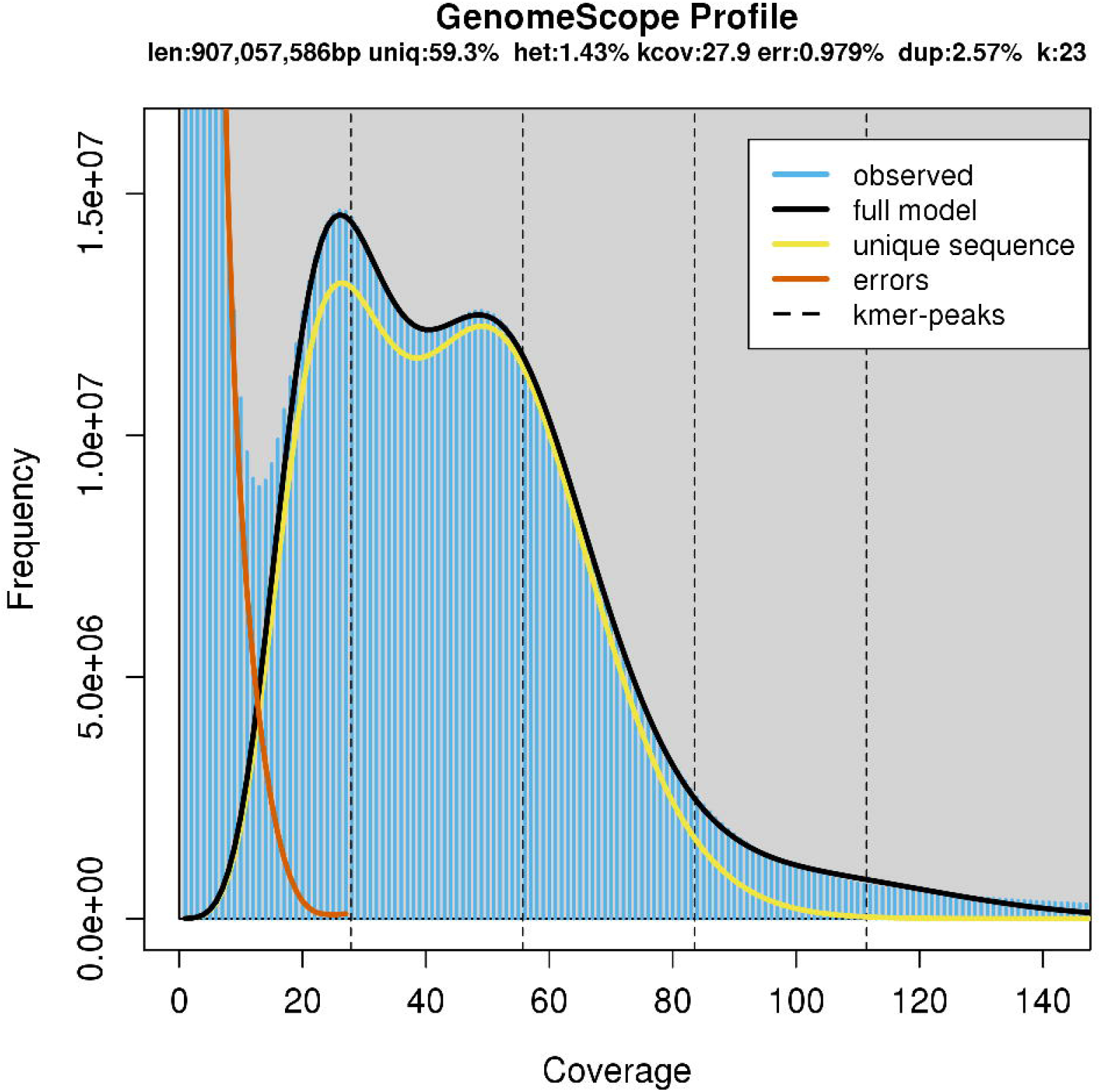
The histogram of the 23-mer depth distribution was plotted in GenomeScope [17] to estimate genome size (907Mb), repeat content (40.7%) and heterozygosity level (1.43%). Two kmer coverage peaks are observed at 28X and 50X.

### *De novo* genome assembly

The *de novo* genome assembly was performed using the paired-end sequence reads from the partitioned library as input for the Supernova assembly algorithm v2.0.0(7fba7b4) (Supernova assembler, RRID:SCR_016756) (10x Genomics, San Francisco, CA, USA) [18]. Two haplotype-resolved genomes, SP_haploid1 (ENA accession ID <u>UOTT01000000</u>) and SP_haploid2 (ENA accession ID <u>UOTU01000000</u>), were assembled with phased scaffolds using the Supernova “mkoutput pseudohap” option. For the assembly process the Supernova run parameters for maximum reads (--maxreads) and barcode fraction (--barfrac) were set for 650M input reads and 80% of barcodes, respectively. Preliminary trials defined an optimal raw coverage of 78-fold, above the 56-fold suggested in the Supernova protocol; this reduced the problem (to some extent) of the complexity of the high repeat content (Table 1). A fraction of the 607.36 million read pairs were used after a quality control step embedded in the Supernova pipeline to remove reads that were not barcoded, not properly paired, or low-quality. Input reads had a 138.5 bp mean length after proprietary 10X barcode trimming and a N50 of 612 per barcode/DNA molecule (Table 1).

**Table 1.** Descriptive metrics, estimated by Supernova, of the input sequence data for the *de novo* genome assembly.

|  |  |
| --- | --- |
| Number of paired reads used | 607.36 M |
| Mean read length after trimming | 138.50 bp |
| Median insert size | 345 bp |
| Weighted mean DNA molecule size | 46.41 Kb |
| N50 reads per barcode | 612 |
| Raw coverage | 78.35 X |
| Effective read coverage | 52.91 X |
| Mean distance between heterozygous SNPs | 197 bp |

Further scaffolding and gap closure procedures were performed with Rails v1.2/Cobbler v0.3 pipeline script [19] to obtain the final consensus genome sequence named SP_G (ENA accession ID GCA_900499035.1) using the parameters anchoring sequence length (*−d* 100) and minimum sequence identity (*−i* 0.95). Three scaffolding and gap closure procedures were performed iteratively with one haplotype of the initial assembly as the assembly *per se*, and previous *de novo* assemblies from Supernova v1.2.2, (315M/100% and 450M/80% reads/barcodes). By closing several gaps within scaffolds and merging other scaffolds into longer and fewer scaffolds (117 259), this procedure resulted into a slightly longer genome size of 949.62 Mb, which slightly deflated the scaffold N50 length to 96.6 Kb (Table 2). The assembly metrics of the three assemblies are described in Table 2 together with a recently published Illumina paired-end assembled sardine genome (UP_Spi) [20]. The total assembly size of our genome (SP_G) is 950 Mb and the UP_Spi is 641 Mb (Table 2). Because the SP_G and UP_Spi assembly sizes are of different orders of magnitude, in addition to N50 we present NG50 values [21] for an estimated genome size of 950 Mb (Table 2). In the SP_G assembly, 905 scaffolds (LG50) represents half of the estimated genome with an NG50 value of 96.6 Kb, in comparison to LG50 of 15 422 and NG50 of 12.6 Kb in the UP_Spi assembly. The ungapped length of the SP_G assembly is 828 Mb. The larger gaps of the SP_G assembly compared to the UP_Spi can be explained by the Supernova being able to estimate gap size based on the bar codes spanning the gaps, i.e. gaps have linkage evidence through the barcodes linking reads to DNA molecules and not solely gaps based on reads pairs [22]. Such gaps are reflected in the large number of N’s per 100kb in our assemblies (Table 2). The number of scaffolds in SP_G is 117 259 (largest 6.843 Mb) and in UP_Spi is 44 627 (largest 0.285 Mb).

**Table 2.** Descriptive metrics of sardine genome assemblies. SP_haploid1/SP_haploid2: haploids genomes (<u>UOTT01000000</u> and <u>UOTU01000000</u>). SP_G: consensus genome (NCBI representative genome assembly, GCA_900499035.1). UP_Spi: Illumina paired-end assembled genome from [20] (GCA_003604335.1). Values for scaffolds equal or larger than 1Kb, 10Kb and 100 Kb are presented in separated rows.

| **Scaffolds** | **Spil_haploid1** | **Spil_haploid2** | **SP_G** | **UP_Spi** |
| --- | --- | --- | --- | --- |
| **Largest** | 6.835 Mb | 6.850 Mb | 6.843 Mb | 0.285 Mb |
| **Number** |  |  |  |  |
| >=100Kb | 874 | 872 | 890 | 309 |
| >= 10Kb | 8 301 | 8 298 | 8 760 | 18 863 |
| >= 1Kb (**total**) | 117 698 | 117 698 | 117 259 | 44 627 |
| **L50 / N50** |  |  |  |  |
| >=100Kb | 135 / 906.0 Kb | 134 / 925.2 Kb | 137 / 899.1 Kb | 130 / 122.5 Kb |
| >= 10Kb | 242 / 572.7 Kb | 242 / 568.2 Kb | 254 / 552.2 Kb | 4 594 / 32.9 Kb |
| >= 1Kb (**total**) | 859 / 102.9 Kb | 860 / 102.7 Kb | 903 / 96.6 Kb | 6 797 / 25.6 Kb |
| **LG50/NG50** | **935 / 87.7 Kb** | **939 / 87.1 Kb** | **905 / 96.6 Kb** | **15 422 / 12.6 Kb** |
| **Assembly size** |  |  |  |  |
| >=100Kb | 469.371 Mb | 468.838 Mb | 473.550 Mb | 39.274 Mb |
| >= 10Kb | 622.165 Mb | 621.688 Mb | 636.491 Mb | 513.719 Mb |
| >= 1Kb (**total**) | 935.548 Mb | 935.082 Mb | 949.618 Mb | 641.169 Mb |
| **GC content** | 43.9 % | 43.9 % | 43.9 % | 44.5 % |
| **N's per 100 Kb** | 12 955 | 12 961 | 12 834 | 169 |

The genome completeness assessment was estimated with Benchmarking Universal Single-copy Orthologs (BUSCO) v3.0.1 (BUSCO, RRID:SCR_015008) [23]. The genome was queried (options -m geno –sp zebrafish) against the “metazoa.odb9” lineage set containing 978 orthologs from sixty-five eukaryotic organisms to assess the coverage of core eukaryotic genes, and against the “actinopterygii.odb9” lineage set containing 4584 orthologs from 20 different ray-finned fish species as the most taxon-specific lineage available for the sardine. Using the metazoan odb9 database, 95.4% of the genome had significant matches: 84.5% were complete genes (76.7% single-copy genes and 9.8% duplicates) and 8.9% were fragmented genes. By contrast, using the actinopterygii odb9 database, 84.2% (76.0% complete genes and 8.2% fragmented) had a match, with 69.3% of genes occurring as single copy and 6.7% as duplicates.

The EMBRIC configurator service [24] was used to create a fish specific checklist (named finfish) for the submission of the sardine genome project to the European Nucleotide Archive (ENA) (European Nucleotide Archive, RRID:SCR_006515) (project accession PRJEB27990).

### Repeat Content

The SP_G consensus assembly was used as a reference genome to build a *de novo* repeat library running RepeatModeler v1.0.11 (RepeatModeler, RRID:SCR_015027) [25] with default parameters. The model obtained from RepeatModeler was used, together with Dfam_consensus database v20171107 [26] and RepBase RepeatMasker Edition library v20170127 [27] to identify repetitive elements and low complexity sequences running RepeatMasker v4.0.7 (RepeatMasker, RRID:SCR_012954) [28]. The analysis carried out revealed that 23.33% of the assembled genome consists of repetitive elements.

### Genome annotation

The Maker v2.31.10 (MAKER, RRID:SCR_005309) [29] pipeline was used iteratively (five times) to annotate the SP_G consensus genome. The annotations generated in each iteration were kept in the succeeding annotation steps and in the final General Feature Format (GFF) file. During the first Maker run the *de novo* transcriptome was mapped to the genome using blastn v2.7.1 (BLASTN, RRID:SCR_001598) [30] (est2genome parameter in Maker). Moreover, the repetitive elements found with RepeatMasker were used in the Maker pipeline. This initial gene models created by Maker were then used to train Hidden Markov Model (HMM) based gene predictors. The preliminary GFF file generated by this first iteration run was used as input to train SNAP v2006-07-28 [31]. Using the scripts provided directly by Maker (maker2zff) and SNAP (fathom, forge and hmm-assembler.pl) an HMM file was created and used as input for the next Maker iteration (snaphmm option in maker configuration file). For the next iteration, the gene-finding software Augustus v3.3 (Augustus, RRID:SCR_008417) [32] was self-trained running BUSCO with the specific parameter (--long), that turn on the Augustus optimization mode for self-training. The resulted predicted species model from Augustus was included in the pipeline in the third Maker run. For the fourth iteration, GeneMark-ES v4.32 (GeneMark, RRID:SCR_011930) [33], a self-training gene prediction software, was executed and the resulting HMM file was integrated into the Maker pipeline. As further evidence for the annotation, in the last run of Maker, the genome was queried using blastx v2.7.1 (BLASTX, RRID:SCR_001653) (protein2genome parameter in Maker), against the deduced proteomes of herring (GCF_000966335.1), (*Clupea harengus*, NCBI:txid7950, Fishbase ID:24) zebrafish (*Danio rerio*, NCBI:txid7955, Fishbase ID:4653) (GCF_000002035.6), blind cave fish (*Astyanax mexicanus*, NCBI:txid7994, Fishbase ID:2740) (GCF_000372685.2), European sardine [20] and all proteins from teleost fishes in the UniProtKB/Swiss-Prot database (UniProtKB, RRID:SCR_004426) [34]. After the five Maker runs the selected 40 777 genes models are the *ab initio* predictions supported by the transcriptome and proteome evidence. Based on the transcriptomic evidence, 12 761 gene models were annotated with untranslated regions (UTR) features, more specifically 9 486 gene models with either 5’ or 3’ UTR and 3 275 gene models with both UTR features. InterProScan v. 5.30 (InterProScan, RRID:SCR_005829) [35] and NCBI blastp v2.8.1 (BLASTP, RRID:SCR_001010) [30] were used to functionally annotate the 40 777 predicted protein coding genes. Thirty-three thousand five hundred and fifty-three (33 553) (82.3%) proteins were successfully annotated using blastp (e-value 1e-05) against the UniProtKB/Swiss-Prot database and another 5 228 were annotated using the NCBI non-redundant protein database (nr). In addition to the above, 37 075 (90.9%) proteins were successfully annotated using InterProScan with all the InterPro v72.0 (InterPro, RRID:SCR_006695) [36] databases: CATH-Gene3D (Gene3D, RRID:SCR_007672), Hamap (HAMAP, RRID:SCR_007701), PANTHER (PANTHER, RRID:SCR_004869), Pfam (Pfam, RRID:SCR_004726), PIRSF (PIRSF, RRID:SCR_003352), PRINTS (PRINTS, RRID:SCR_003412), ProDom (ProDom, RRID:SCR_006969), ProSite Patterns (PROSITE, RRID:SCR_003457), ProSite Profiles, SFLD (Structure-function linkage database, RRID:SCR_001375), SMART (SMART, RRID:SCR_005026), SUPERFAMILY (SUPERFAMILY, RRID:SCR_007952), and TIGRFAM (JCVI TIGRFAMS, RRID:SCR_005493). In total, 38 880 (95.3%) of the predicted proteins received a functional annotation. The annotated genome assembly is published [37] in the wiki-style annotation portal ORCAE [38].

OrthoFinder v2.2.7 [39] was used to identify paralogy and orthology in our Swiss-prot annotated deduced proteome and in the deduced proteomes from herring, blind cave fish and zebrafish. The resulting orthogroups were plotted using jvenn (jVenn, RRID:SCR_016343) [40] (Figure 3), where paralagous (two or more genes) and singletons were identified within species specific orthogroups. The deduced sardine proteome has 3 413 paralogous groups containing 11 406 genes, of which 31 are sardine specific orthogroups. The amount of sardine singletons (9 856) can be partially due to fragmented predicted genes, but can reflect also some evolutionary divergence which requires further study to understand the biological relevance. In total, 25 560 orthogroups containing at least a single protein were identified in sardine, of which 12 958 ortholgroups are common to all four fish species. Within the Clupeidae, the sardine and the herring share 14 780 orthogroups with 922 family-specific orthogroups.

**Figure 3.**
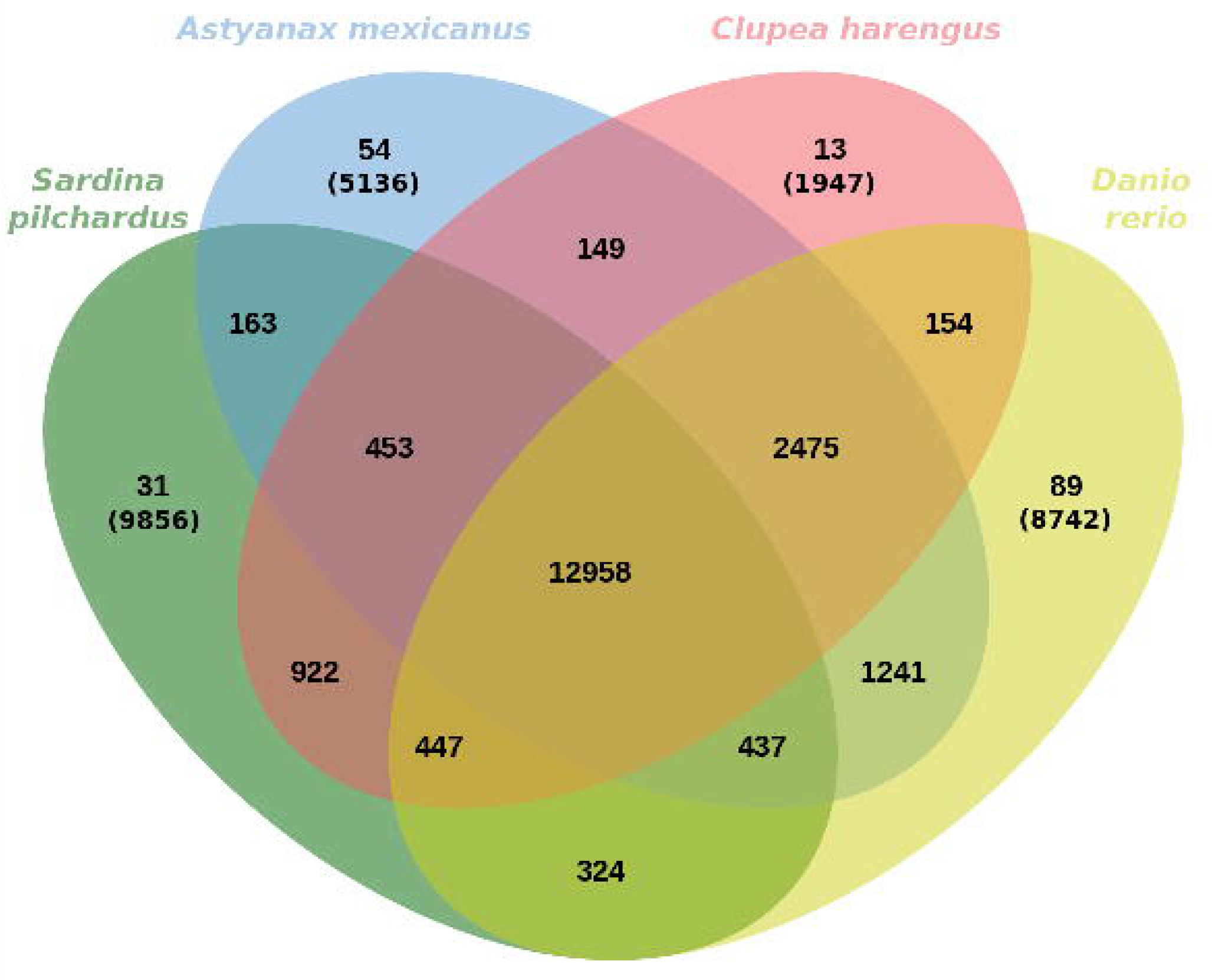
Optimal maximum likelihood tree (-ln likelihood: 146565.6438) under a best-fitting JTT+Γ_4_+F substitution model of 97 concatenated proteins. Maximum likelihood bootstrap support values are given below or to the right of nodes. Scale bar represents mean numbers of substitutions per site. The Actinopterygii ingroup was rooted to two outgroup taxa, namely *Petromyzon marinus* (lamprey) and *Latimeria chalumnae* (coelacanth) (not shown).

### Variant calling between phased alleles

FASTQ files were processed using the 10x Genomics LongRanger v2.2.2 pipeline [41] with a maximum input limit of one thousand scaffolds, defined as reference genome, and representing about half of the genome size (488.5 Mb). The LongRanger pipeline was run with default settings, with the exception of vcmode to define the Genome Analysis Toolkit (GATK) v4.0.3.0 (GATK, RRID:SCR_001876) [42] as the variant caller and the somatic parameters. The longest phase block was 2.86 Mb and the N50 phase block was 0.476 Mb.

Single nucleotide polymorphisms (SNP’s) were furthered filtered to obtain only phased and heterozygous SNP’s between the two alleles with a coverage higher than 10-fold using VCFtools v0.1.16 (VCFtools, RRID:SCR_001235). A VCF file was obtained containing 2 369 617 filtered SNPs (Additional file 1) resulting in a mean distance between heterozygous phased SNPs of 206 bp. Similar results were obtained in the Supernova input report estimation (Table 1) of mean distance between heterozygous SNPs in the whole genome of 197 bp. This high SNP heterozygosity (1/206), observed solely in the comparison of the phased alleles, is higher than the average nucleotide diversity of the previously reported marine fish of wild populations: 1/390 in yellow drum [43], 1/309 in herring [44], 1/435 in coelacanth [45], 1/500 in cod [46] and 1/700 in stickleback [47].

### *De novo* transcriptome assembly

The 596 million paired-end raw transcriptomic reads were edited for contamination (e.g. adapters) using TrimGalore v0.4.5 wrapper tool (TrimGalore, RRID:SCR_016946) [15], low-quality base trimming with Cutadapt v1.15 (cutadapt, RRID:SCR_011841) [48] and the output overall quality reports of the edited reads with FastQC v0.11.5 (FastQC, RRID:SCR_014583) [49].

The 553 million edited paired-end reads were *de novo* assembled as a multi-tissue assembly using Trinity v2.5.1 (Trinity, RRID:SCR_013048) [50] with a minimum contig length of 200 bp, 50x coverage read depth normalization, and RF strand-specific read orientation. The same parameters were used for each of the 11 tissue specific *de novo* assemblies. The genome and transcriptome assemblies were conducted on the Portuguese National Distributed Computing Infrastructure [49].

The twelve *de novo* transcriptome assemblies (Table 3) were each quality assessed using TransRate v1.0.3 [51] with read evidence for assembly optimization, by specifying the contigs fasta file and respective left and right edited reads to be mapped. The multi-tissue assembly with all reads resulted in an assembled transcriptome of 170 478 transcript contigs following the TransRate step. Functional annotation was performed using the Trinotate v3.1.1 pipeline [24] and integrated into a SQLite database. All annotations were based on the best deduced open reading frame (ORF) obtained with the Transdecoder v1.03 [51]. Of the 170 478 transcripts contigs, 27 078 (16%) were inferred to ORF protein sequences. Query of the UniProtKB/Swiss-Prot (e-value cutoff of 1e-5) database via blastx v2.7.1 of total contigs resulted in 43 458 (26%) annotated transcripts. The ORFs were queried against UniProtKB/Swiss-Prot (e-value cutoff of 1e-5) via blastp v2.7.1 and PFAM using hmmscan (HMMER v3.1b2) (Hmmer, RRID:SCR_005305) [52] resulting in 19 705 (73% of ORF) and 16 538 (61% of ORF) UniProtKB/Swiss-Prot and PFAM annotated contigs respectively. The full annotation report with further functional annotation, such as signal peptides, transmembrane regions, eggnog, Kyoto Encyclopedia of Genes and Genomes (KEGG) (KEGG, RRID:SCR_012773), and Gene Ontology annotation (Gene Ontology, RRID:SCR_002811) are listed in tabular format in Additional file 2.

**Table 3.** Summary statistics of transcriptome data for the eleven tissues.

| Tissue | Paired raw reads | Contigs | CDS deduced | SwissProt annotated | Accession number |
| --- | --- | --- | --- | --- | --- |
| Gill/Branchial Arch | 29 783 994 | 62 526 | 29.3% | 38.6% | ERS2629269 |
| Liver | 33 479 471 | 53 104 | 29.7% | 40.1% | ERS2629273 |
| Spleen | 25 634 530 | 66 419 | 31.6% | 40.4% | ERS2629276 |
| Ovary | 22 241 327 | 42 521 | 38.1% | 42.5% | ERS2629270 |
| Midgut | 28 016 117 | 75 782 | 31.0% | 39.5% | ERS2629274 |
| White Muscle | 24 409 160 | 49 266 | 35.4% | 44.8% | ERS2629277 |
| Red Muscle | 30 653 774 | 55 873 | 30.3% | 42.1% | ERS2629275 |
| Kidney | 27 861 879 | 59 495 | 30.8% | 37.3% | ERS2629272 |
| Head Kidney | 25 280 960 | 65 888 | 32.2% | 38.4% | ERS2629271 |
| Brain/Pituitary | 24 467 352 | 75 620 | 24.5% | 37.1% | ERS2629267 |
| Caudal Fin<br>(Skin/Cartilage/Bone) | 26 342 097 | 64 832 | 23.9% | 38.0% | ERS2629268 |
| All Tissues | 298 170 661 | 170 478 | 15.9% | 25.5% | ERS2629362 |

### Ray-finned fish phylogeny

We conducted a phylogenetic analysis of ray-finned fish (Actinopterygii) taxa based on 17 fish species. The sardine protein data set used in the phylogenetic analysis was obtained by querying the deduced proteins from our sardine genome against the one-to-one orthologous cluster dataset (106 proteins from 17 species) obtained from [20].

For the query, gene models were constructed for each protein with hmmbuild (HMMER v3.1b2) [53] using default options and the orthologous genes from the deduced sardine proteome were searched using hmmsearch (HMMER) with an e-value cuttoff of 10e-3. The best protein hits, as indicated by the bitscores, were aligned to the original protein sequence alignments using hmmalign (HMMER) with default options. Gapped and poorly aligned sites were identified by Gblocks v0.91b (Gblocks, RRID:SCR_015945) [54] using default options and removed using p4 v1.3.0 [55]. Protein alignment statistics were calculated, and the proteins concatenated into a single alignment using novel scripts in p4. Of the 106 fish proteins alignments, 97 contained sites which were considered correctly aligned by the Gblocks analysis; statistics for these alignments are presented in Table S1 (Additional file 3). The concatenated sequence alignment of the 97 proteins contained 14 515 sites without gaps of which 7 391 were constant, 7 123 variable, and 3 879 parsimony informative.

The best-fitting empirical protein model of the concatenated data was evaluated using ModelFinder [56] in IQ-TREE v1.6.7.1 [57]. The best-fitting empirical substitution model was estimated to be the JTT model [58] with a discrete gamma-distribution of among-site rate variation (4 categories) and empirical composition frequencies (typical notation: JTT+Γ_4_+F).

Optimal maximum likelihood tree searches (100 replicates) and bootstrap analyses (300 replicates) were conducted using RAxML v8.2.12 (RAxML, RRID:SCR_006086) [59] with the best-fitting model. The optimal maximum likelihood tree (-ln likelihood: 146565.6438) is presented in Figure 4 with bootstrap support values given at nodes, and is rooted to the outgroups *Petromyzon marinus* (lamprey) and *Latimeria chalumnae* (coelacanth).

**Figure 4.**
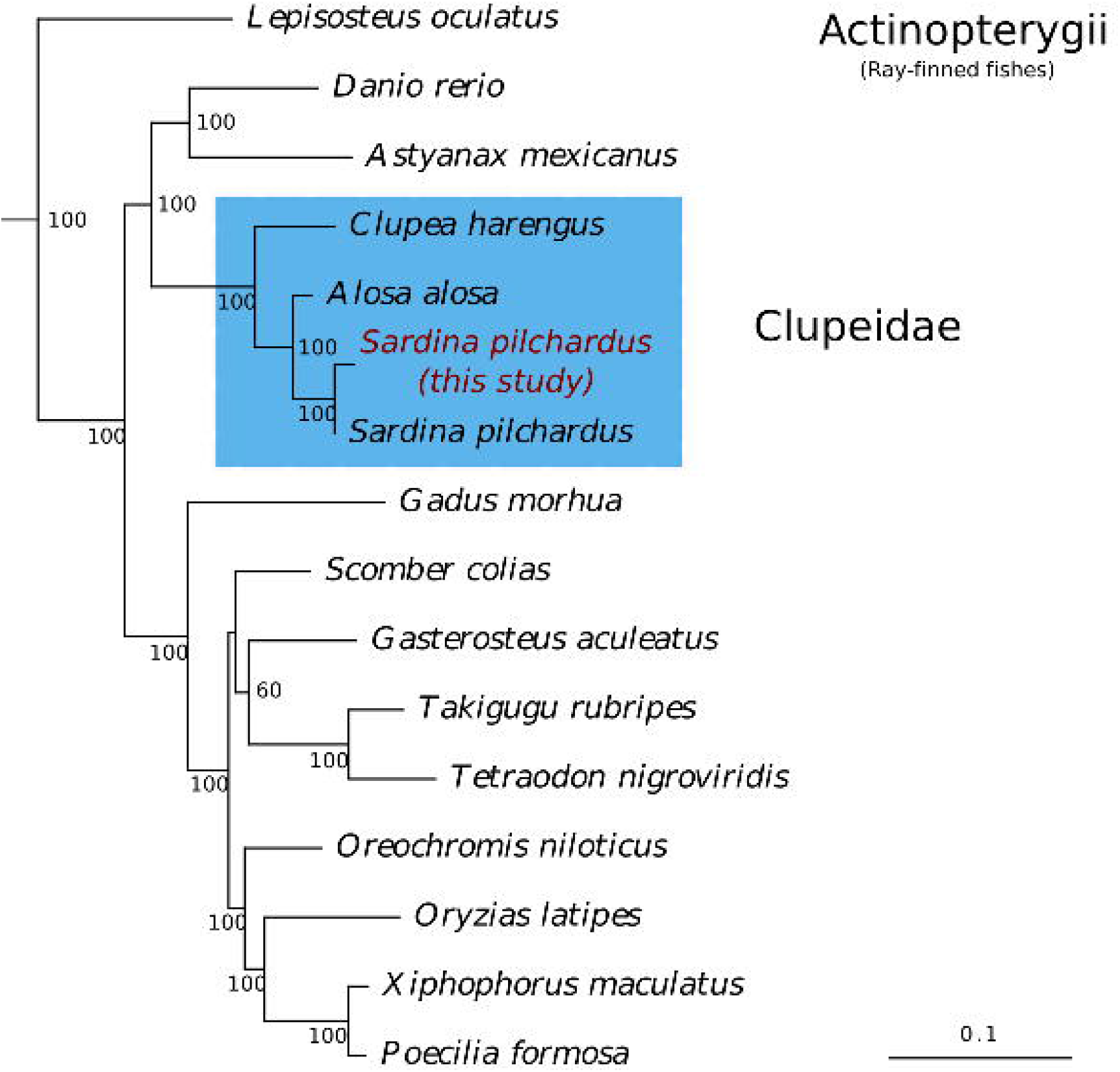
Venn diagram representing paralogous and orthologous groups between sardine, blind cave fish, zebrafish, and herring obtained with OrthoFinder and plotted with Jvenn [40]. Orthogroups of singleton genes are showed in parenthesis.

## Conclusion

Despite the sardine genome having a high level of repeats and heterozygosity, factors which pose a challenge to *de novo* genome assembly, a more than adequate draft genome was obtained with the 10X Genomics linked-reads (Chromium) technology. The Chromium technology’s ability to tag and cluster the reads to individual DNA molecules has proven advantages for scaffolding, just as long reads technologies such as Nanopore and Pacific Biosciences, but with high coverage and low error rates. The advantage of linked-reads for *de novo* genomic assemblies is evident in comparison to typical short read data, especially in the case of wild species with highly heterozygous genomes, where the latter often result in many uncaptured genomic regions and with a lower scaffolding yield due to repeated content.

The high degree of heterozygosity identified here in the sardine genome illustrates l future problems for monitoring sardine populations using low-resolution genetic data. However, the phased SNPs obtained in this study can be used to initiate the development of a SNP genetic panel for population monitoring, with SNPs representative of haplotype blocks, allowing insights into the patterns of linkage disequilibrium and the structure of haplotype blocks across populations.

The genomic and transcriptomic resources reported here are important tools for future studies to understand sardine response at the levels of physiology, population genetics and ecology of the causal factors responsible for the recruitment and collapse of the sardine stock in Iberian Atlantic coast. Besides the commercial interest, the sardine plays a crucial role at a key trophic level by bridging energy from the primary producers to the top predators in the marine ecosystem. Therefore, disruption of the sardine population equilibrium is likely to reverberate throughout the food chain via a trophic cascade. Consequently, these genomic and genetic resources are the prerequisites needed to develop tools to monitor the population status of the sardine and thereby provide an important bio-monitoring system for the health of the marine environment.

## Availability of the supporting data

Raw data, assembled transcriptomes, and assembled genomes are available at the European Bioinformatics Institute ENA archive with the project accession PRJEB27990. The annotated genome assembly is published in the wiki-style annotation portal ORCAE [37].

## Acknowledgements

This research was supported by national funds from FCT – Foundation for Science and Technology through project UID/Multi/04326/2016 and by FCT and FEDER under projects 22153-01/SAICT/2016 (to INCD), ALG-01-0145-FEDER-022121 and ALG-01-0145-FEDER-022231; and co-funds from MAR2020 operational programme of the European Maritime and Fisheries Fund (project SARDINOMICS MAR-01.04.02-FEAMP-0024). The EMBRIC configurator service received funding from the European Union’s Horizon 2020 research and innovation programme under grant agreement No 654008. The authors acknowledge Pedro Guerreiro for providing the sardine samples.

## Additional files

**Additional file 1.** Heterozygous SNPs identified in the phased haploid blocks listed in a VCF file format.

**Additional file 2.** Annotation of all tissues transcriptome assembly in a tabular format.

**Additional file 3.** Sequence alignment statistics of the 97 proteins concatenated for the phylogenetics analyses.

