## Additional file 3 for "A haplotype-resolved draft genome of the European sardine (*Sardina pilchardus*)"

Table S1. Sequence alignment statistics of the 97 proteins concatenated for the phylogenetics analyses. The concatenated sequence alignment of the 97 proteins contained 14,515 sites without gaps of which 7391 were constant, 7123 variable, and 3879 parsimony informative.

|  | **Alignment sites** | | | | |
| --- | --- | --- | --- | --- | --- |
| **Gene Name** | **total** | **constant** | **variable** | **ambiguous** | **pars_infor** |
| cmtr1 | 139 | 64 | 75 | 0 | 36 |
| psmd6 | 384 | 295 | 89 | 0 | 40 |
| nme4 | 120 | 18 | 102 | 0 | 52 |
| ddx18 | 379 | 281 | 98 | 0 | 62 |
| atp6v0d1 | 213 | 185 | 28 | 0 | 5 |
| trappc3 | 99 | 73 | 26 | 0 | 18 |
| tmem64 | 20 | 2 | 18 | 0 | 10 |
| dera | 276 | 155 | 121 | 9 | 85 |
| wdr45 | 199 | 130 | 69 | 0 | 27 |
| tsn | 165 | 68 | 97 | 1 | 31 |
| glo1 | 154 | 100 | 54 | 0 | 28 |
| jmjd7 | 85 | 37 | 48 | 0 | 32 |
| naa20 | 128 | 92 | 36 | 0 | 8 |
| mrpl46 | 53 | 13 | 40 | 0 | 27 |
| tfb2m | 176 | 17 | 159 | 0 | 104 |
| cox11 | 183 | 118 | 65 | 0 | 38 |
| med10 | 79 | 46 | 33 | 0 | 13 |
| cactin | 221 | 150 | 71 | 0 | 35 |
| rab20 | 118 | 50 | 68 | 81 | 35 |
| hectd3 | 425 | 74 | 351 | 17 | 119 |
| nanp | 175 | 47 | 128 | 0 | 95 |
| gba2 | 349 | 143 | 206 | 0 | 113 |
| mccc2 | 119 | 92 | 27 | 0 | 13 |
| hmces | 45 | 15 | 30 | 0 | 15 |
| suclg1 | 271 | 219 | 52 | 0 | 38 |
| coq9 | 190 | 46 | 144 | 0 | 68 |
| ube2g2 | 86 | 74 | 12 | 0 | 5 |
| yipf4 | 58 | 47 | 11 | 0 | 5 |
| zgc | 81 | 29 | 52 | 0 | 25 |
| glod4 | 213 | 53 | 160 | 0 | 70 |
| si | 89 | 36 | 53 | 0 | 31 |
| fam120b | 64 | 18 | 46 | 0 | 32 |
| tbc1d7 | 173 | 64 | 109 | 0 | 72 |
| washc3 | 56 | 28 | 28 | 0 | 16 |
| phb | 255 | 178 | 77 | 0 | 33 |
| slc35b2 | 210 | 144 | 66 | 0 | 40 |
| ddost | 414 | 220 | 194 | 0 | 76 |
| sharpin | 68 | 22 | 46 | 1 | 27 |
| ndufa6 | 122 | 46 | 76 | 0 | 42 |
| gemin2 | 16 | 1 | 15 | 0 | 6 |
| gemin8 | 100 | 38 | 62 | 18 | 42 |
| sspn | 40 | 7 | 33 | 0 | 24 |
| atp6v1d | 243 | 175 | 68 | 0 | 39 |
| rxylt1 | 80 | 27 | 53 | 0 | 37 |
| mrpl51 | 64 | 24 | 40 | 0 | 31 |
| bloc1s4 | 81 | 29 | 52 | 0 | 24 |
| apex1 | 277 | 152 | 125 | 0 | 72 |
| cgrrf1 | 83 | 12 | 71 | 0 | 57 |
| sra1 | 72 | 18 | 54 | 0 | 32 |
| enoph1 | 149 | 47 | 102 | 0 | 52 |
| slc38a9 | 238 | 106 | 132 | 0 | 71 |
| dcaf12 | 135 | 87 | 48 | 0 | 20 |
| hspa13 | 17 | 11 | 6 | 2 | 3 |
| mrps10 | 113 | 43 | 70 | 0 | 40 |
| psmb3 | 147 | 105 | 42 | 0 | 8 |
| mlst8 | 271 | 213 | 58 | 0 | 17 |
| timm29 | 179 | 39 | 140 | 0 | 106 |
| pak1ip1 | 138 | 32 | 106 | 0 | 71 |
| exoc8 | 570 | 335 | 235 | 0 | 134 |
| setd4 | 39 | 17 | 22 | 0 | 11 |
| pex2 | 51 | 21 | 30 | 51 | 22 |
| FAM109B | 128 | 54 | 74 | 0 | 33 |
| txndc5 | 304 | 107 | 197 | 0 | 109 |
| phax | 98 | 32 | 66 | 0 | 45 |
| TIMM21 | 24 | 6 | 18 | 0 | 4 |
| ndufv2 | 119 | 43 | 76 | 1 | 35 |
| gtf2h3 | 173 | 108 | 65 | 0 | 34 |
| ttc36 | 170 | 72 | 98 | 0 | 67 |
| rnaseh2a | 48 | 21 | 27 | 0 | 18 |
| cmpk2 | 135 | 38 | 97 | 0 | 56 |
| tsku | 138 | 49 | 89 | 0 | 56 |
| abhd13 | 279 | 130 | 149 | 0 | 67 |
| mrpl32 | 30 | 13 | 17 | 0 | 13 |
| acot13 | 111 | 47 | 64 | 0 | 32 |
| gclm | 191 | 78 | 113 | 0 | 68 |
| gatad1 | 49 | 30 | 19 | 0 | 11 |
| clpp | 173 | 130 | 43 | 0 | 22 |
| ndufaf1 | 137 | 45 | 92 | 0 | 61 |
| mtfr1 | 32 | 10 | 22 | 0 | 11 |
| cdkn2aipnl | 50 | 22 | 28 | 0 | 20 |
| crnkl1 | 642 | 445 | 197 | 0 | 126 |
| ap2s1 | 134 | 131 | 3 | 0 | 2 |
| serpine2 | 138 | 19 | 119 | 0 | 76 |
| med29 | 127 | 80 | 47 | 0 | 21 |
| sap30l | 96 | 84 | 12 | 0 | 4 |
| scnm1 | 44 | 16 | 28 | 0 | 17 |
| mrps25 | 131 | 53 | 78 | 0 | 46 |
| ube2s | 107 | 30 | 77 | 0 | 13 |
| zcchc17 | 48 | 16 | 32 | 0 | 19 |
| srp19 | 118 | 50 | 68 | 0 | 35 |
| llph | 90 | 40 | 50 | 0 | 37 |
| slc35b4 | 157 | 77 | 80 | 0 | 34 |
| hypk | 66 | 47 | 19 | 0 | 8 |
| mcee | 91 | 56 | 35 | 0 | 20 |
| glrx5 | 88 | 56 | 32 | 0 | 20 |
| ecsit | 189 | 78 | 111 | 0 | 67 |
| gosr2 | 102 | 50 | 52 | 0 | 32 |
| Concatenated | 14514 | 7391 | 7123 | - | 3879 |

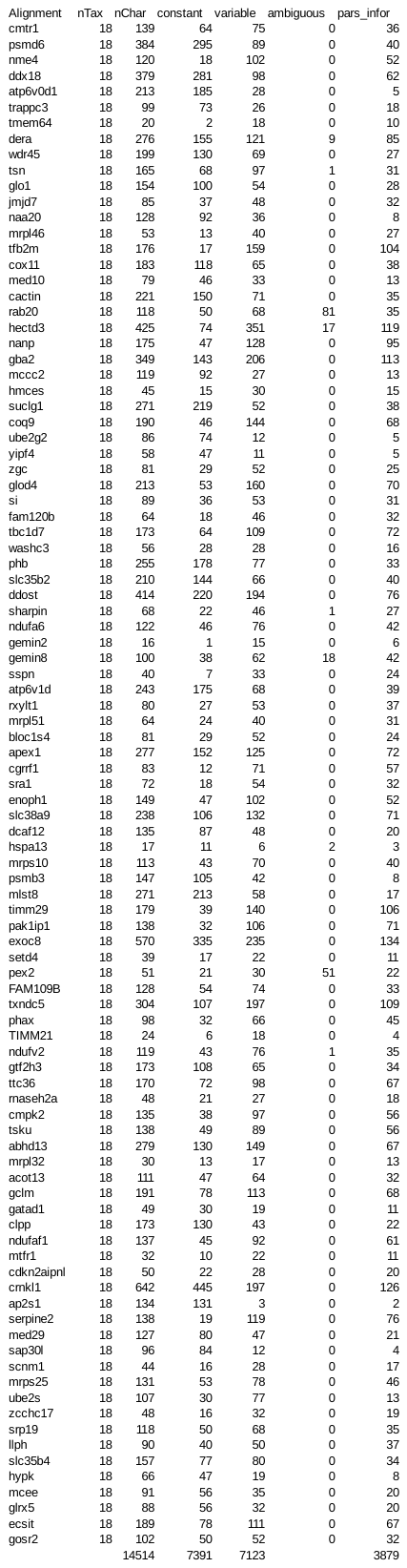
